# Hypoxia impedes differentiation of cranial neural crest cells into derivatives relevant for craniofacial development

**DOI:** 10.1101/2024.09.10.610634

**Authors:** Theresa Schmid, Gabriele Rodrian, Alexander Kohler, Michael Wegner, Lina Gölz, Matthias Weider

## Abstract

Orofacial clefts are the second-most prevalent congenital malformation. Risk factors are multifactorial and include genetic components but also environmental factors. One environmental factor is hypoxia during pregnancy, caused for instance by tobacco smoking, medication or living at high altitudes. Knowledge about the molecular link between hypoxia and orofacial clefts is at large. We here show that hypoxia has only modest effects on proliferating cranial neural crest cells, but dramatically influences their differentiation potential. We detected massive perturbations in their differentiation to chondrocytes, osteoblasts and smooth muscle cells. The transcriptional induction of the majority of regulated genes during each of these processes was grossly impaired by hypoxic conditions, as evidenced by genome-wide transcriptomic analyses. Bioinformatic analyses pointed to cytoskeletal organization and amino acid metabolism as two main processes compromised during all three differentiation pathways, and several orofacial cleft risk genes were among the genes with impaired induction during hypoxia. Our analyses reveal a drastic influence of hypoxia on the differentiation potential of cranial neural crest cells as a possible source for the occurrence of orofacial clefts.

## INTRODUCTION

Orofacial clefts (OFC) represent the second-most prevalent congenital malformation (Leslie and Marazita, 2013; Mossey et al., 2009). They manifest as cleft lip with or without cleft palate (CL/P) or as isolated cleft palate (CPO). Depending on ethnic and geographic variation, CL/P rates range from 1 in 3,000 to 1 in 450 births (Mossey et al., 2009). CPO prevails with more pronounced variation and even higher maximum frequency, ranging from 1 in 8,000 to 1 in 400 (Mossey et al., 2009). OFC can occur as part of a syndrome or as non-syndromic forms. OFC patients need long-lasting multi-disciplinary treatment, as they suffer from various problems affecting dentition, feeding, hearing and speech. Even after successful surgical intervention OFC often pose a life-long psychological burden on patients. Craniofacial development is one of the most complex developmental events during embryogenesis and depends on a substantial contribution of a transient, pluripotent stem cell-like cell population called cranial neural crest cells (CNCC) (Nichols, 1981; Roth et al., 2021; Weston and Thiery, 2015). CNCC are specified from neuroectodermal cells in the direct vicinity of the neural plate (Weston and Thiery, 2015). With the completion of neurulation, i.e. the closure of the neural tube, neural crest cells undergo epithelial-to-mesenchymal transition and delaminate. These ectomesenchymal CNCC are highly migratory. They proliferate massively during migration into the pharyngeal arches – bilateral outgrowths along the pharyngeal area (Weston and Thiery, 2015). Later in craniofacial development, CNCC differentiate into a multitude of tissues and specialized cells such as cartilage, bone, vessel smooth musculature, adipose tissue, (para)sympathetic neurons, peripheral glia, pigment cells, as well as dentin-forming odontoblasts in the pulp of teeth (Achilleos and Trainor, 2012). Indeed, most craniofacial structures are derived from CNCC (Roth et al., 2021). Proper craniofacial development therefore depends largely on CNCC, but also on ectodermal epithelial cells and the interactions of both cell types.

OFC arise when growth, cell differentiation processes or fusion events of craniofacial tissues occur incorrectly. The fusion of medial and lateral nasal prominences with maxillary prominences, forming the upper lip, is the starting point of palate development (Bush and Jiang, 2012; Dixon, Michael J. et al., 2011). For formation of the secondary palate, the palatal shelves that are derived from the maxillary prominences and which are located in a vertical position next to the tongue, are elevated above the tongue, grow horizontally towards each other until they gain contact and fuse. Fusion failure of the medial or lateral nasal prominences and maxillary prominences gives rise to cleft lip, whereas fusion failure of both palatal shelves leads to cleft palate. Fusion failure of anterior structures is often accompanied with clefts of posterior tissues. Thus, cleft lip frequently goes along with cleft palate (Mangold et al., 2017).

The etiology of OFC is multifactorial and involves genetic predispositions. Additionally, environmental factors play a crucial role in pathogenesis of non-syndromic OFC (nsOFC) (Babai and Irving, 2023; Mossey et al., 2009). A well-known risk factor is alcohol consumption during pregnancy (DeRoo et al., 2008). Vitamin shortage seems to be a risk factor, too. It could for instance be demonstrated that folic acid supplements reduce the risk of OFC (Babai and Irving, 2023; Mossey et al., 2009). Maternal smoking is considered as the strongest risk factor (Burg et al., 2016; Mossey et al., 2009). However, it is not easy to distinguish between the effects of teratogens contained in smoke and the effect of fetal hypoxia caused by carbon monoxide (Longo, 1976; Mossey et al., 2009; Socol et al., 1982). Nevertheless, other findings indicate that hypoxia is a causative factor for OFC. The anticonvulsant phenytoin is a broadly known example for medication that evokes OFC when taken during pregnancy (Czeizel, 1976). It was suggested that phenytoin intake results in fetal hypoxia due to slowed heart rates (Webster et al., 2006). This assumption is strengthened by the finding that hyperoxia during embryonic development reduces the incidence of OFC in mice exposed to phenytoin (Millicovsky and Johnston, 1981). In a clinical study, acardiac monozygotic twins displayed a high incidence of OFC of 50%, likely due to low oxygenation (Jones et al., 2008). Moreover, zebrafish embryos exposed to hypoxia display a gap in the anterior edge of the ethmoid plate, the equivalent to OFC in mammals (Küchler et al., 2018).

Final proof that hypoxia can evoke OFC came from a mouse study in which pregnant mice were kept at 10% O_2_ during the time of fusion of nasal and maxillary prominences (Millicovsky and Johnston, 1981). Fetuses from mothers of the hypoxic group displayed shortly before birth a substantially higher occurrence of CL/P compared to fetuses from mothers under normal atmospheric oxygen levels of 21%. In humans, living at high altitudes is associated with a significantly higher relative risk for OFC, most likely because high altitudes cause chronic hypobaric hypoxia. It has been shown that pregnancy at high altitudes (>2,000 m) is associated with cleft lip and other craniofacial defects (Castilla et al., 1999) as well as with nsCL/P (Poletta et al., 2007). Indeed, newborns at high altitudes display a reduced arterial oxygen saturation during their first hours of life (Li et al., 2023). Hence, according to official statistics, Bolivia is the country with the highest reported prevalence rate for CL/P and a similar rate seems to exist for the Mongolian population of Tibet (Mossey and Catilla, 2003; Mossey et al., 2009).

Mechanistic understanding of how hypoxia evokes OFC is lacking, although such knowledge is central for prevention of OFC in risk pregnancies. We therefore set out to analyze the molecular changes in CNCC when exposed to hypoxic conditions.

## RESULTS

### Hypoxia has only mild impact on proliferating CNCC

In order to address how hypoxia might influence craniofacial development and lead to OFC, we chose a CNCC cell culture model. We used the O9-1 cell line, because it is a well characterized cellular model of CNCC and the cells can be differentiated to three tissues that are relevant for palatogenesis, i.e. chondrocytes, osteoblasts and smooth muscle cells (Ishii et al., 2012). To study the effects of hypoxia, we first kept cells under proliferating conditions and compared normal atmospheric oxygen levels (21%) in the CO_2_ incubator with 0.5% oxygen in a hypoxia chamber. As expected, we detected induction of all tested genes that are characteristic for hypoxia such as *Gapdh* (Graven et al., 1994), *Glut1* (Gleadle and Ratcliffe, 1997), *Pdk1* (Kim et al., 2006) and *Bnip3* (Guo et al., 2001) (Figs. 1A-D). We examined the proliferation of O9-1 cells for 72 h. At 24 h and 48 h cell numbers of hypoxic and normoxic cells had increased equally (Figs. 1E,F). Only after 72 h in hypoxia, we could detect a small, but statistically not significant decrease of relative cell numbers. Accordingly, life/dead staining revealed identical percentages of dead cells in hypoxia as compared to normoxia at 24 h and 48 h, and only a small, but statistically not significant increase of cell death at 72 h (Figs. 1G,H).

**Fig. 1.**
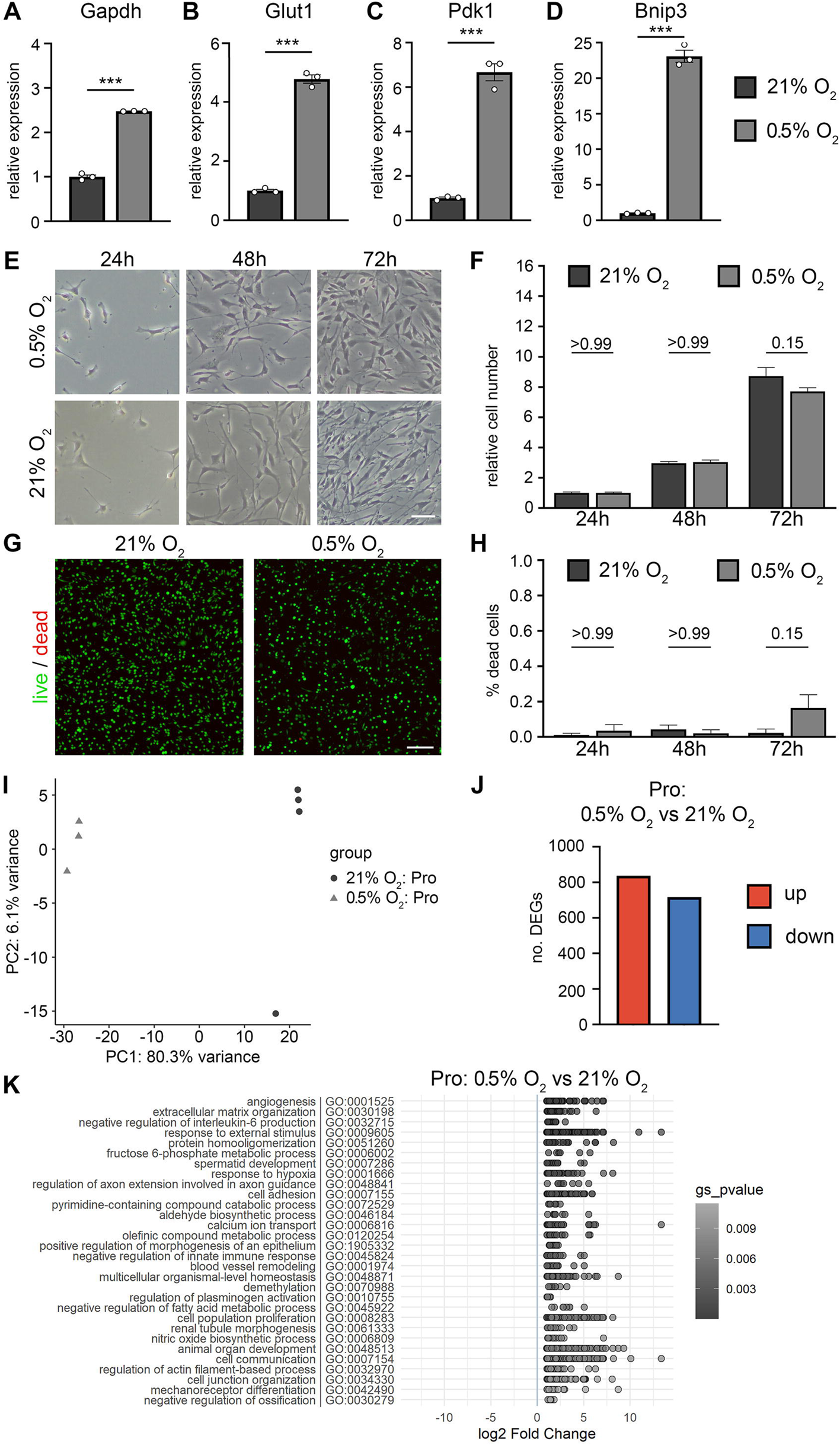
Hypoxia has only mild impact on proliferating CNCC. **A – D:** Relative expression of hypoxic marker genes *Gapdh* (a), *Glut1* (b), *Pdk1* (c) and *Bnip3* (d) in proliferating O9-1 cranial neural crest cells as determined by qRT-PCR at normal atmospheric oxygen levels (21% O_2_) and at hypoxia (0.5% O_2_); n = 3; ***0.0001 ≤ p < 0.01. **E:** Representative photographs of proliferating O9-1 cells grown at 21% O_2_ or 0.5% O_2_ for 24 h, 48 and 72 h. **F:** Quantification of proliferating O9-1 cell numbers grown for 24 h, 48 h or 72 h at 21% O_2_ or 0.5% O_2_. Values were referenced to arithmetic means at T = 24 h and are shown ±SEM; n = 3; p-values for pairwise comparisons of the different O_2_ concentrations are indicated above columns. **G:** Representative photographs of live/dead staining of proliferating O9-1 cells grown for 72 h at 21% O_2_ or 0.5% O_2_. Green calcein fluorescence depicts life cells while red propidium iodide fluorescence depicts dead cells. **H:** Quantification of dead O9-1 cells kept for 24 h, 48 h or 72 h in proliferation medium at 21% O_2_ or 0.5% O_2_ as determined as percentage of total cells ±SEM; n = 3; p-values for pairwise comparisons of the different O_2_ concentrations are indicated above columns. **I:** Principal component analysis (PCA) on the gene expression value (FPKM) of proliferating O9-1 cells (Pro) under normoxic (21% O_2_, n=4) and hypoxic conditions (0.5% O_2_, n=3). **J:** Number of upregulated (red) and downregulated genes (blue) of proliferating O9-1 cells under hypoxic (0.5% O_2_) compared to normoxic (21% O_2_) conditions (|log2-fold| ≥ 1, p-value ≤ 0.05, mean of normalized counts ≥ 100). **K:** Gene ontology analysis of upregulated genes in hypoxic compared to normoxic conditions of proliferating O9-1 cells. Size bars: 20 µm (e) and 100 µm (g).

We asked which genes are deregulated under hypoxic conditions in O9-1 cells and performed bulk RNA-seq experiments. As expected, normoxic and hypoxic samples clustered separately in principle component analysis (PCA) (Fig. 1i). We found that 837 genes were upregulated, while 717 genes were downregulated in hypoxia (Fig. 1J, |log2(FoldChange)| ≥ 1, mean count ≥ 100, p-value ≤ 0.05). Gene Ontology (GO) analysis of induced genes expectedly yielded terms associated with hypoxia such as angiogenesis, response to hypoxia, blood vessel remodeling, nitric oxide biosynthesis and IL-6 production (Fig. 1K). Additionally, GO terms related to cell adhesion, cell junctions and cytoskeleton were enriched in the induced genes.

We were astonished to see only rather mild effects on proliferation and apoptosis, because 0.5% oxygen represents a severe hypoxic condition. We therefore speculated that other processes must be responsible for the emergence of hypoxia-evoked orofacial clefts and decided to analyze the differentiation of O9-1 cells to the three available derivatives chondrocytes, osteoblasts and smooth muscle cells.

### Hypoxia impedes differentiation of CNCC to chondrocytes

We differentiated O9-1 cells to chondrocytes according to established protocols (Ishii et al., 2012) under normoxia (21% O_2_) and in a hypoxic chamber (0.5% O_2_). Induction of hypoxia-related genes *Gapdh, Glut1, Pdk1* and *Bnip3* proved that the cells grown under chondrogenic conditions are affected by hypoxia as well (Fig. 2A). We analyzed the success of the differentiation regime by qRT-PCR of the collagen genes *Col2a1* and *Col11a2* and compared proliferative conditions and chondrogenic differentiation with 21% oxygen and 0.5% oxygen. Intriguingly, compared to normoxia, induction of both genes was hardly detectable under hypoxic conditions and there was no statistical significance when compared to proliferating cells (Figs. 2B,C). This massive inhibition of differentiation-related induction under hypoxia prompted us to investigate the ongoing process by bulk RNA-seq. The samples clustered according to their growth condition in a PCA plot, with chondrogenic differentiated cells set clearly apart from proliferating cells and hypoxic cells separated from the respective normoxic conditions (Fig. 2D). The finding that hypoxic chondrocytes still were in close proximity to normoxic chondrocytes suggests that differentiation to chondrocytes is – although impaired – per se still possible. We analyzed the number of deregulated genes (DEGs) and found 1931 genes upregulated and 1506 genes downregulated during chondrogenic differentiation in normoxia (Fig. 2E). Differentiation under hypoxic conditions gave similar numbers of deregulated genes (Fig. 2F): 1557 genes were upregulated and 1405 genes were downregulated when compared to proliferation under hypoxic conditions. When chondrogenic differentiation under hypoxic conditions was compared to normoxic conditions, only 169 genes were induced and 251 genes were downregulated (Fig. 2G). This lead us to the hypothesis that chondrogenic differentiation per se is still possible under hypoxic conditions, even though the extent of differentiation is reduced dramatically (cf. Figs. 2B,C).

**Fig. 2.**
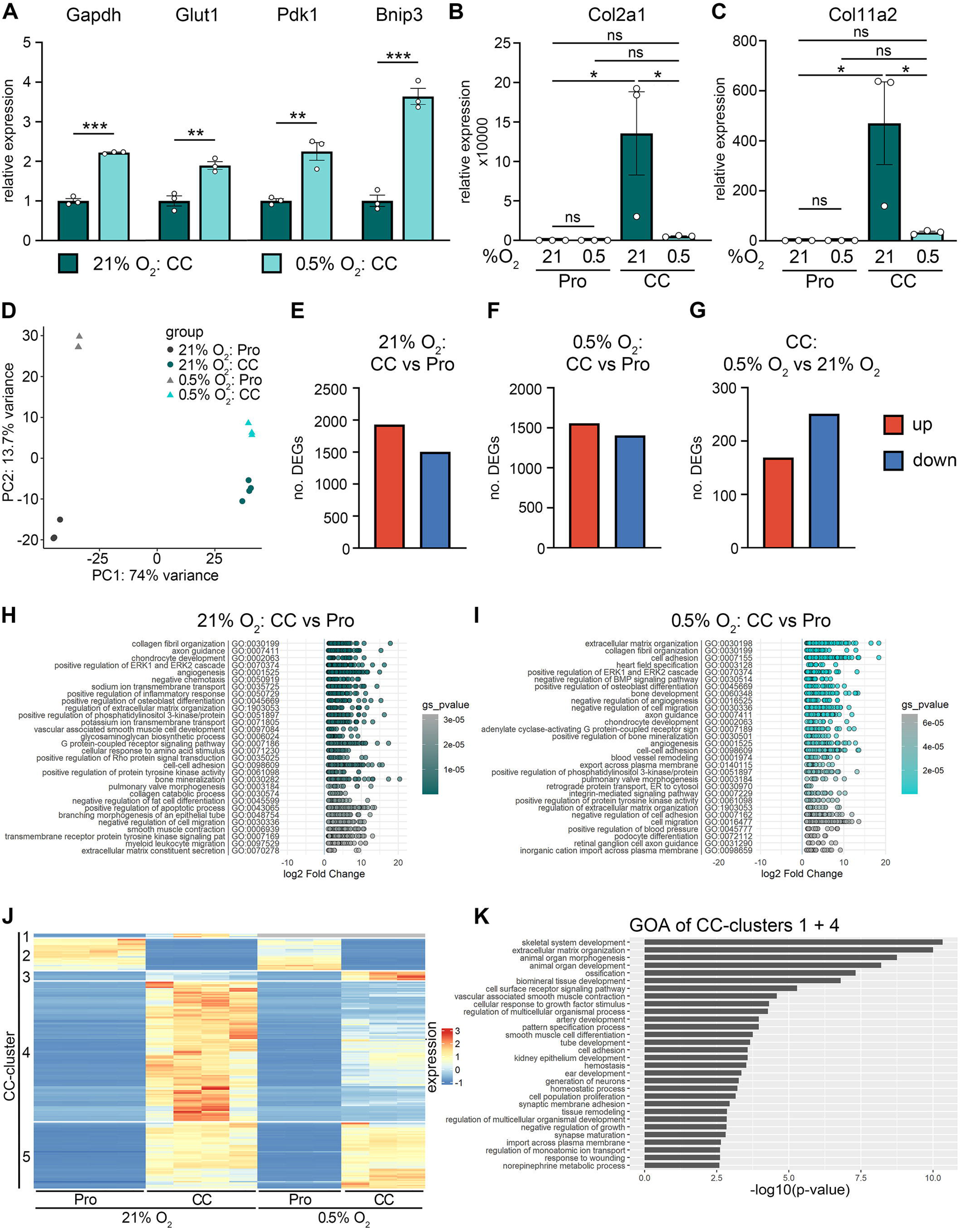
Hypoxia impedes differentiation of CNCC to chondrocytes. **A:** Relative expression of hypoxic marker genes *Gapdh, Glut1, Pdk1* and *Bnip3* in O9-1 cranial neural crest cells differentiated to chondrocytes as determined by qRT-PCR at normal atmospheric oxygen levels (21% O_2_) and at hypoxia (0.5% O_2_). Graphs show arithmetic mean ±SEM. Values for chondrocyte-differentiated cells at 21% O_2_ were arbitrarily set to 1; n = 3. **B, C:** Relative expression of chondrocyte marker genes *Col2a1* (b) and *Col11a2* (c) in O9-1 cells differentiated to chondrocytes (CC) compared to proliferating O9-1 cells (Pro) as determined by qRT-PCR at normal atmospheric oxygen levels (21% O_2_) and at hypoxia (0.5% O_2_). Graphs show arithmetic mean ±SEM. Values for proliferating cells at 21% O_2_ were arbitrarily set to 1; n = 3. **D:** Principal component analysis (PCA) on the gene expression values (FPKM) of proliferating (Pro) and chondrocyte-differentiated (CC) O9-1 cells under normoxic (n = 4 each) and hypoxic conditions (n = 3 each). **E:** Number of upregulated (red) and downregulated genes (blue) of chondrocyte-differentiated vs. proliferating O9-1 cells under normoxic condition (|log2-fold| ≥ 1, p-value ≤ 0.05, mean of normalized counts ≥ 100). **F:** Number of upregulated (red) and downregulated genes (blue) of chondrocyte-differentiated vs. proliferating O9-1 cells under hypoxic condition (|log2-fold| ≥ 1, p-value ≤ 0.05, mean of normalized counts ≥ 100). **G:** Number of upregulated (red) and downregulated genes (blue) of chondrocyte-differentiated O9-1 cells cultivated in hypoxic (0.5% O_2_) vs. normoxic (21% O_2_) conditions (|log2-fold| ≥ 1, p-value ≤ 0.05, mean of normalized counts ≥ 100). **H, I:** Gene ontology analysis of upregulated genes in chondrocyte-differentiated O9-1 cells compared to proliferating O9-1 cells under normoxic conditions (21% O_2_, h) and under hypoxic conditions (0.5% O_2_, i). **J:** Heatmap of top 200 differentially expressed genes of chondrocyte-differentiated compared to proliferating O9-1 cells under normoxic conditions. Scaled normalized counts for proliferating (Pro) and chondrocyte-differentiated (CC) O9-1 cells at normoxic (21% O_2_) and hypoxic conditions (0.5% O_2_) are shown. Clusters 1 and 4 were analyzed further. **K:** Gene ontology analysis of genes in clusters 1 and 4 from Fig. 2j. *0.01 ≤ p < 0.05, **0.001 ≤ p < 0.01, ***0.0001 ≤ p < 0.01.

We asked whether certain molecular processes are specifically altered during differentiation under hypoxic conditions. Therefore, we performed GO analysis of induced genes during chondrogenic differentiation under normoxia (Fig. 2H) or hypoxia (Fig. 2I). In principle, both conditions yielded similar enriched GO terms. Among the 30 most significant GO terms of each condition were expected categories such as collagen fibril organization, chondrocyte development and regulation of extracellular matrix organization. Moreover, terms related to osteoblast differentiation and bone development were also enriched in both settings, probably due to an overlap of gene expression in bone and chondrocyte differentiation. Marked differences appeared in terms related to cell adhesion: while in normoxia only cell-cell adhesion was enriched, we additionally detected cell adhesion, integrin-mediated signaling pathway and negative regulation of cell adhesion among the significantly enriched processes under hypoxia.

We speculated that hampered induction during chondrogenic differentiation as detected for *Col2a1* and *Col11a2* (cf. Fig. 2B;C) is true for the majority of induced genes. Therefore, we clustered the 200 mostly deregulated genes under normoxia using hclust in R (Fig. 2J). We could identify five distinct clusters: (CC-cluster 1 (CC1)) induced under 21% O_2_ but not expressed under 0.5% O_2_; (CC2) strongly expressed in proliferating conditions and repressed during chondrogenic differentiation under 21% O_2_, while weaker expressed (but also repressed) under 0.5% O_2_; (CC3) mildly induced during differentiation under 21% O_2_, strongly induced during differentiation under 0.5% O_2_; (CC4) strongly induced during chondrogenic differentiation under 21% O_2_ and only mildly induced during differentiation under 0.5% O_2_; (CC5) similarly induced under normoxia and hypoxia (gene lists are presented in supplementary table 1). Indeed, the majority of genes was only weakly induced or not expressed in hypoxic conditions (115 of 200 deregulated genes, clusters 1&4). We like to introduce the term “hypoxia-attenuated genes” for genes that are induced during differentiation in normoxic conditions, but only faintly induced or not expressed in hypoxia.

We sought to identify biological processes important for chondrogenic differentiation that are hampered under hypoxia. To this aim, we performed GO analysis of the hypoxia-attenuated genes during chondrogenic differentiation (i.e. genes of CC clusters 1 and 4) (Fig. 2K). Interestingly, we did not find terms related to chondrocytes (except extracellular matrix organization) but rather yielded terms related to osteoblasts such as skeletal system development, ossification and biomineral tissue development and diverse biological processes that were not related. Thus, chondrogenic differentiation per se is still possible under hypoxic conditions, but the induction of relevant genes is drastically compromised, affecting a variety of biological processes.

### Hypoxia hampers differentiation of CNCC to osteoblasts

We asked if differentiation of CNCC to osteoblasts is similarly impaired and performed differentiation of O9-1 cells to osteoblasts (Ishii et al., 2012) under 21% O_2_ and 0.5% O_2_. All tested hypoxia markers were again induced, as proven by qRT-PCR (Fig. 3A). Similar to the situation during chondrogenic differentiation, we could not detect induction of the osteoblast marker genes *Msx1* and *Runx2* under hypoxic conditions (Figs. 3B,C). We performed bulk RNA-seq and PCA showed distinct clustering of samples according to growth conditions (Fig. 3D). Osteoblast differentiation resulted in 2238 induced and repressed genes each under 21% O_2_ (Fig. 3E), and 1736 induced and 1453 repressed genes under 0.5% O_2_ (Fig. 3F). When we compared differentiation to osteoblasts under both conditions, we detected 980 genes that were expressed at higher levels under hypoxia and 801 that were expressed at lower levels than under normoxia (Fig. 3G).

**Fig. 3.**
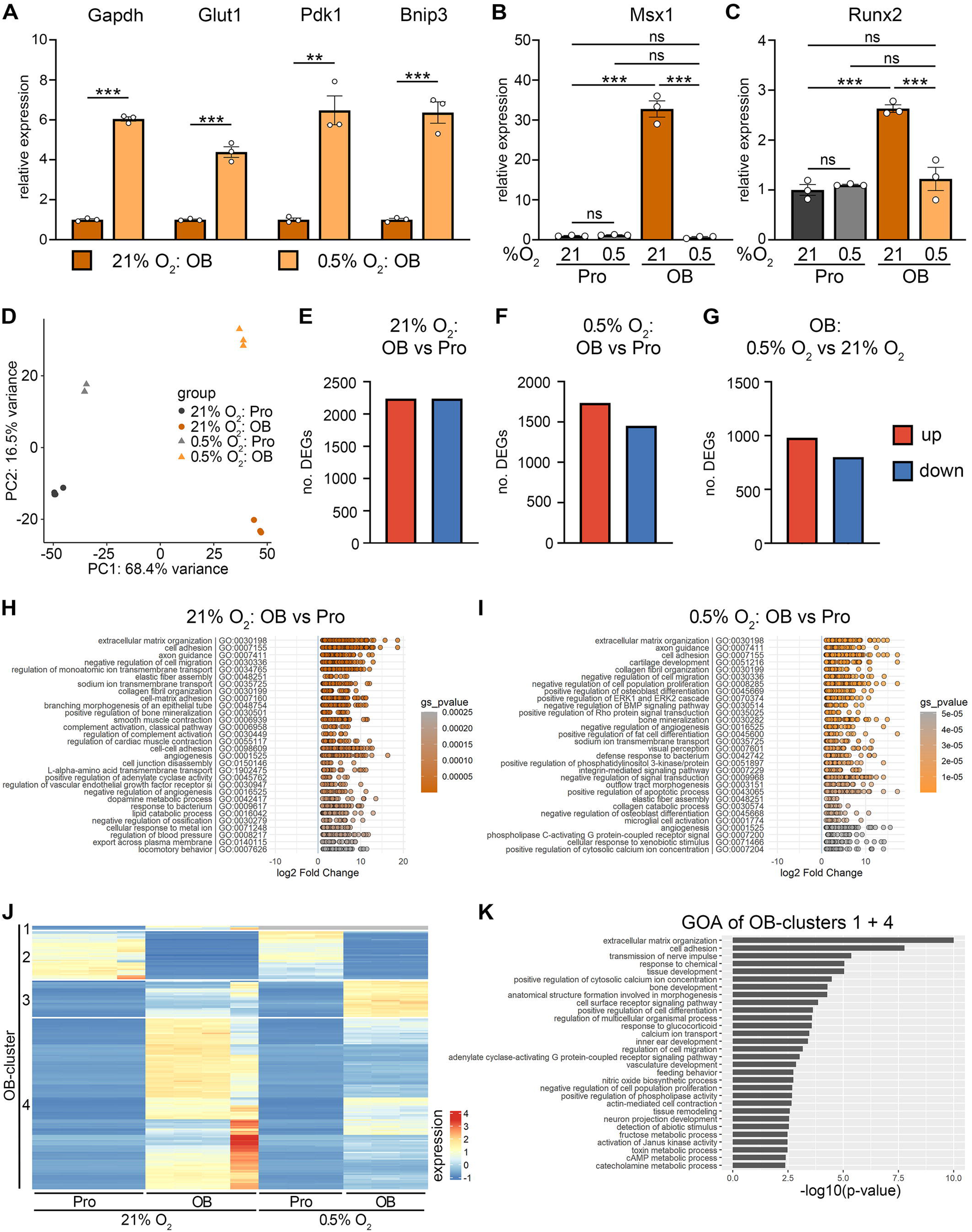
Hypoxia impedes differentiation of CNCC to osteoblasts. **A:** Relative expression of hypoxic marker genes *Gapdh, Glut1, Pdk1* and *Bnip3* in O9-1 cranial neural crest cells differentiated to osteoblasts as determined by qRT-PCR at normal atmospheric oxygen levels (21% O_2_) and at hypoxia (0.5% O_2_). Graphs show arithmetic mean ±SEM. Values for osteoblast-differentiated cells at 21% O_2_ were arbitrarily set to 1; n = 3. **B, C:** Relative expression of osteoblast marker genes *Msx1* (b) and *Runx2* (c) in O9-1 cells differentiated to osteoblasts (OB) compared to proliferating O9-1 cells (Pro) as determined by qRT-PCR at normal atmospheric oxygen levels (21% O_2_) and at hypoxia (0.5% O_2_). Graphs show arithmetic mean ±SEM. Values for proliferating cells at 21% O_2_ were arbitrarily set to 1; n = 3. **D:** Principal component analysis (PCA) on the gene expression values (FPKM) of proliferating (Pro) and osteoblast-differentiated (OB) O9-1 cells under normoxic (n = 4 each) and hypoxic conditions (n = 3 each). **E:** Number of upregulated (red) and downregulated genes (blue) of osteoblast-differentiated vs. proliferating O9-1 cells under normoxic condition (|log2-fold| ≥ 1, p-value ≤ 0.05, mean of normalized counts ≥ 100). **F:** Number of upregulated (red) and downregulated genes (blue) of osteoblast-differentiated vs. proliferating O9-1 cells under hypoxic condition (|log2-fold| ≥ 1, p-value ≤ 0.05, mean of normalized counts ≥ 100). **G:** Number of upregulated (red) and downregulated genes (blue) of osteoblast-differentiated O9-1 cells cultivated in hypoxic (0.5% O_2_) vs. normoxic (21% O_2_) conditions (|log2-fold| ≥ 1, p-value ≤ 0.05, mean of normalized counts ≥ 100). **H, I:** Gene ontology analysis of upregulated genes in osteoblast-differentiated O9-1 cells compared to proliferating O9-1 cells under normoxic conditions (21% O_2_, h) and under hypoxic conditions (0.5% O_2_, i). **J:** Heatmap of top 200 differentially expressed genes of osteoblast-differentiated compared to proliferating O9-1 cells under normoxic conditions. Scaled normalized counts for proliferating (Pro) and osteoblast-differentiated (OB) O9-1 cells at normoxic (21% O_2_) and hypoxic conditions (0.5% O_2_) are shown. Clusters 1 and 4 were analyzed further. **K:** Gene ontology analysis of genes in clusters 1 and 4 from Fig. 3j. **0.001 ≤ p < 0.01, ***0.0001 ≤ p < 0.01.

GO enrichment analysis (Figs. 3H,I) still yielded terms related to osteoblast differentiation when cells were differentiated under hypoxic conditions, for example positive regulation of osteoblast differentiation, negative regulation of BMP signaling pathway and bone mineralization (Fig. 3I). Thus, osteoblast differentiation also seems possible under hypoxic conditions in principle. We compared expression of the 200 most deregulated genes during normoxic osteoblast differentiation by hierarchical clustering using R function hclust, which helped to identify four clusters (Fig. 3J): (OB-cluster 1 (OB1)) induced under 21% O_2_ but not expressed under 0.5% O_2_; (OB2) expressed in proliferating conditions and repressed during chondrogenic differentiation under 21% O_2_ and under 0.5% O_2_; (OB3) mildly induced during differentiation under 21% O_2_, strongly induced during differentiation under 0.5% O_2_; (OB4) strongly induced during chondrogenic differentiation under 21% O_2_ and only mildly induced during differentiation under 0.5% O_2_ (gene lists are presented in supplementary table 1). Resembling the situation for the osteoblast marker genes *Msx1* and *Runx2* (cf. Figs. 3B,C), the majority of genes induced during normoxic osteoblast differentiation was only very weakly induced or not expressed under hypoxic conditions (135 of 200 deregulated genes, OB clusters 1&4).

In order to identify biological processes that are mostly affected by hypoxia, we performed GO analysis of the hypoxia-attenuated genes (Fig. 3K). We did not find striking GO terms that could point to a larger impairment of differentiation to osteoblasts such as bone metabolism, ossification or similar. Rather, terms belonged to different processes such as extracellular matrix organization, cell adhesion and calcium ion transport or regulation. Osteoblast differentiation therefore is drastically hampered under hypoxia, obviously with no specific process particularly affected but rather a broad spectrum of differentiation genes severely impaired in their induction.

### Hypoxia hinders differentiation of CNCC to smooth muscle cells

The obvious question arose if the same would be true for the third possible differentiation paradigm related to craniofacial development, i.e. smooth muscle differentiation. Again, we performed differentiation of O9-1 cells to smooth muscle cells (Ishii et al., 2012) under 21% O_2_ and 0.5% O_2_. As expected, the cells responded to 0.5% O_2_ with the induction of all tested hypoxia marker genes also under differentiation to smooth muscle cells (Fig. 4A). Strikingly, the induction of both tested marker genes for smooth muscles *Acta2* and *Tagln* was severely diminished (Figs. 4B,C). Bulk RNA-seq and PCA yielded specific clusters for proliferating and differentiated cells under normoxia and hypoxia (Fig. 4D). 1946 genes were induced and 1905 were repressed during smooth muscle differentiation at 21% O_2_ (Fig. 4E). We found similar numbers of 2038 induced and 1891 repressed genes during differentiation at 0.5% O_2_ (Fig. 4F). When comparing smooth muscle differentiation at hypoxia with normoxia, we identified high numbers of 2330 induced and 2551 downregulated genes (Fig. 4G).

**Fig. 4.**
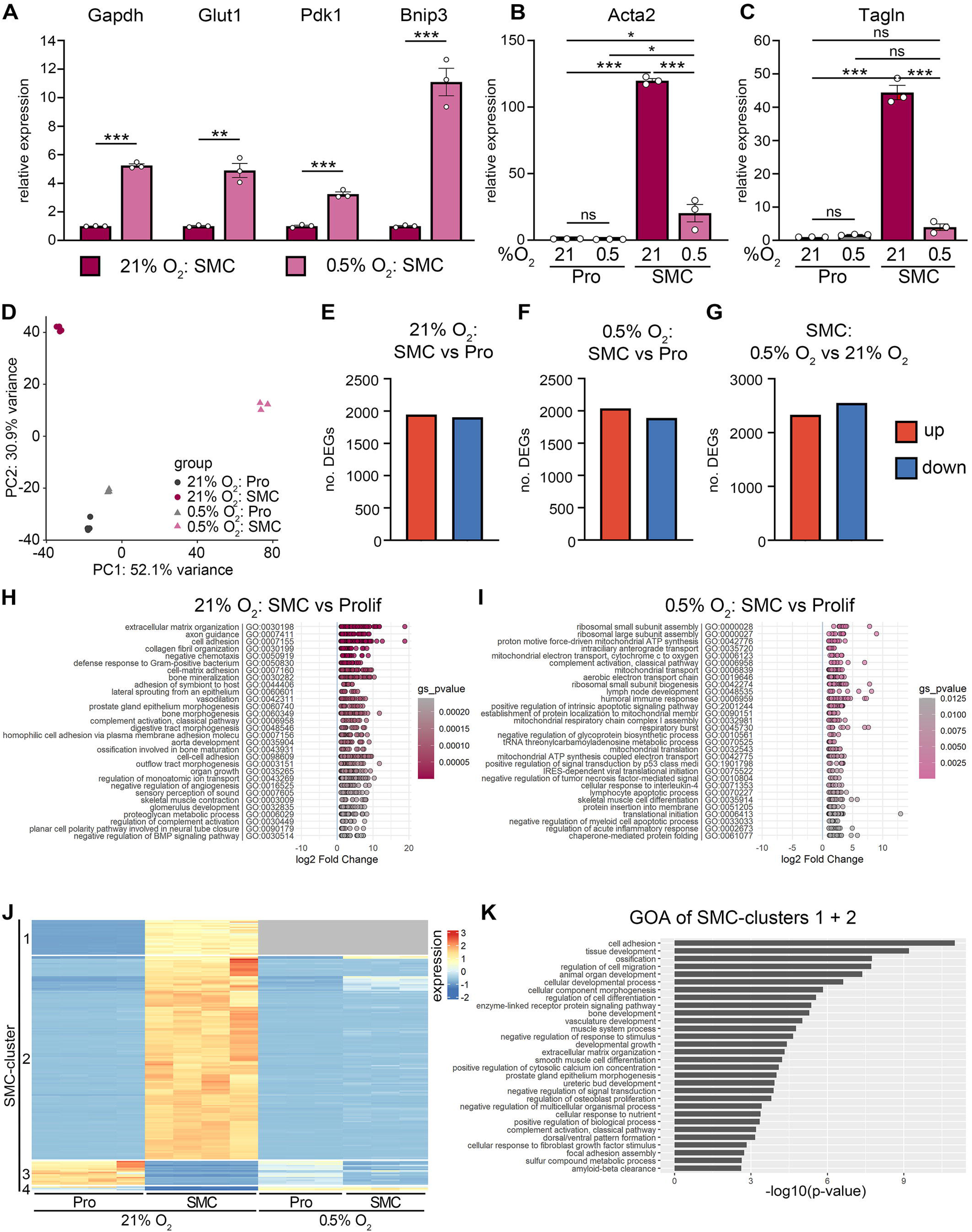
Hypoxia impedes differentiation of CNCC to smooth muscle cells. **A:** Relative expression of hypoxic marker genes *Gapdh, Glut1, Pdk1* and *Bnip3* in O9-1 cranial neural crest cells differentiated to smooth muscle cells as determined by qRT-PCR at normal atmospheric oxygen levels (21% O_2_) and at hypoxia (0.5% O_2_). Graphs show arithmetic mean ±SEM. Values for smooth muscle cell-differentiated cells at 21% O_2_ were arbitrarily set to 1; n = 3. **B, C:** Relative expression of smooth muscle cell marker genes *Acta2* (b) and *Tagln* (c) in O9-1 cells differentiated to smooth muscle cells (SMC) compared to proliferating O9-1 cells (Pro) as determined by qRT-PCR at normal atmospheric oxygen levels (21% O_2_) and at hypoxia (0.5% O_2_). Graphs show arithmetic mean ±SEM. Values for proliferating cells at 21% O_2_ were arbitrarily set to 1; n = 3. **D:** Principal component analysis (PCA) on the gene expression values (FPKM) of proliferating (Pro) and smooth muscle cell-differentiated (SMC) O9-1 cells under normoxic (n = 4 each) and hypoxic conditions (n = 3 each). **E:** Number of upregulated (red) and downregulated genes (blue) of smooth muscle cell-differentiated vs. proliferating O9-1 cells under normoxic condition (|log2-fold| ≥ 1, p-value ≤ 0.05, mean of normalized counts ≥ 100). **F:** Number of upregulated (red) and downregulated genes (blue) of smooth muscle cell-differentiated vs. proliferating O9-1 cells under hypoxic condition (|log2-fold| ≥ 1, p-value ≤ 0.05, mean of normalized counts ≥ 100). **G:** Number of upregulated (red) and downregulated genes (blue) of smooth muscle cell-differentiated O9-1 cells cultivated in hypoxic (0.5% O_2_) vs. normoxic (21% O_2_) conditions (|log2-fold| ≥ 1, p-value ≤ 0.05, mean of normalized counts ≥ 100). **H, I:** Gene ontology analysis of upregulated genes in smooth muscle cell-differentiated O9-1 cells compared to proliferating O9-1 cells under normoxic conditions (21% O_2_, h) and under hypoxic conditions (0.5% O_2_, i). **J:** Heatmap of top 200 differentially expressed genes of smooth muscle cell-differentiated compared to proliferating O9-1 cells under normoxic conditions. Scaled normalized counts for proliferating (Pro) and smooth muscle cell-differentiated (SMC) O9-1 cells at normoxic (21% O_2_) and hypoxic conditions (0.5% O_2_) are shown. Clusters 1 and 2 were analyzed further. **K:** Gene ontology analysis of genes in clusters 1 and 2 from Fig. 4j. *0.01 ≤ p < 0.05, **0.001 ≤ p < 0.01, ***0.0001 ≤ p < 0.01.

GO term analysis of genes induced at normoxia generated terms related to blood vessel and muscle development and function such as lateral sprouting, vasodilation, aorta development, outflow tract morphogenesis, negative regulation of angiogenesis and skeletal muscle contraction (Fig. 4H). In contrast, at hypoxic conditions only 1 of the 30 most statistically significant GO terms belonged to these categories (i.e. skeletal muscle cell differentiation, Fig. 4I). Instead, the largest category of GO terms was related to mitochondrial electron chain and aerobic ATP production (proton motive force-driven ATP synthesis, mitochondrial electron transport - cytochrome c to oxygen, aerobic electron transport chain and respiratory burst), probably because the smooth muscle cells were undergoing a known hypoxia-induced shift to mitochondrial respiration (Akagi et al., 2023). Another large category of GO terms was related to ribosome assembly and translation (ribosomal small subunit assembly, ribosomal large subunit assembly, ribosomal small subunit biogenesis and translational initiation).

Using hierarchical clustering of the 200 top-deregulated genes during normoxic smooth muscle differentiation, we determined 4 clusters (Fig. 4J): (SMC-cluster 1 (SMC1)) induced under 21% O_2_ but not expressed under 0.5% O_2_; (SMC2) induced under 21% O_2_ but only very faintly induced under 0.5% O_2_; (SMC3) expressed in proliferating conditions and repressed during smooth muscle differentiation under 21% O_2_, weakly expressed at 0.5% O_2_ and repressed during differentiation 0.5% O_2_; (SMC4) repressed during differentiation at 21% O_2_ and induced at 0.5% O_2_ (gene lists are presented in supplementary table 1). This showed that most of the genes induced during normoxic differentiation were only very faintly induced under hypoxic conditions or not expressed at all (179 of 200). GO term analysis of hypoxia-attenuated genes (SMC clusters 1&2) pointed to impaired differentiation to smooth muscle cells (vasculature development, muscle system development and smooth muscle cell differentiation) but also to compromised induction of genes more generally related to differentiation (tissue development, ossification, regulation of cell differentiation, bone development and extracellular matrix organization) (Fig. 4K). CNCC thus presumably still kept their potential to differentiate to smooth muscle cells, although induction of relevant genes was faint (cf. Figs. 4B,C).

### Hypoxia interferes with cytoskeletal organization and amino acid metabolism during differentiation

In order to identify common disturbances that might be caused by hypoxia in all three differentiation paradigms, we compared hypoxia-attenuated genes of chondrocyte differentiation, osteoblast differentiation and smooth muscle differentiation (Fig. 5A). There was an overlap of 22 genes for all three conditions, corresponding to 19.1% of genes not induced in chondrocytes, 16.3% not in osteoblasts and 12.3% not in smooth muscle cells. We performed gene ontology analysis of those commonly induced genes that failed to be induced in hypoxia during differentiation (Fig. 5B) and mostly detected terms related to differentiation and development in rather unrelated terms (Fig. 5C). Only two categories could be found in more terms, which were related to cytoskeleton on the one hand (actin-mediated cell contraction, regulation of actin filament bundle assembly, cell adhesion) and amino acid metabolism on the other (cysteine metabolic process, serine family amino acid catabolic process, L-arginine transmembrane transport, cellular modified amino acid catabolic process).

**Fig. 5.**
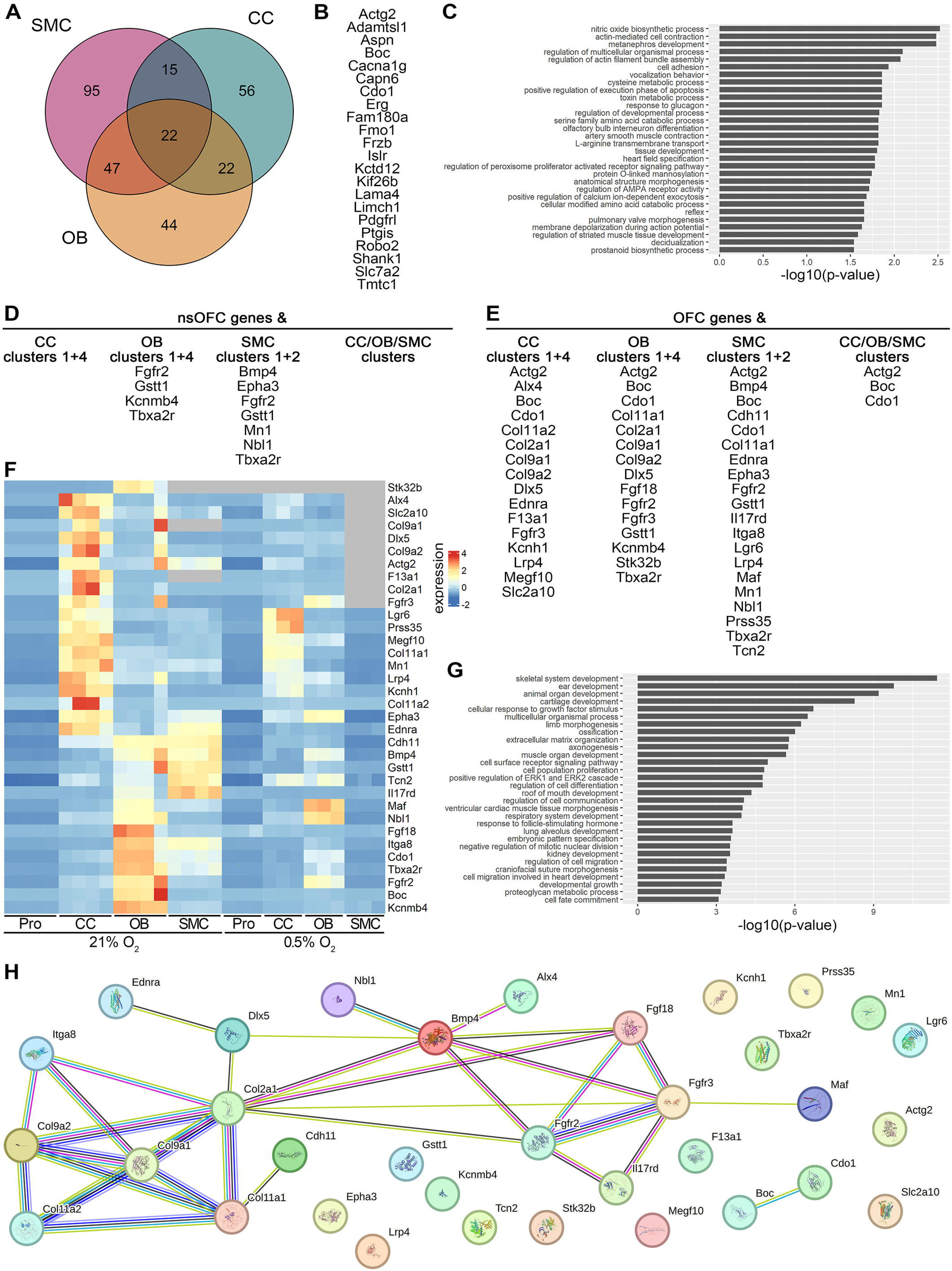
Hypoxia constrains induction of several OFC risk genes and limits induction of some commonly induced genes during differentiation to chondrocytes, osteoblasts and smooth muscle cells. **A:** Venn diagram of genes from clusters 1 and 4 from Fig. 2j and Fig. 3j and clusters 1 and 2 from Fig. 4j depicting hypoxia-constrained induction of genes during differentiation to chondrocytes, osteoblasts or smooth muscle cells and of genes shared between different or all three pathways. B: List of common genes with hypoxia-constrained induction, i.e. genes that are weakly induced, not induced or not expressed in hypoxia during differentiation to chondrocytes, osteoblasts as well as smooth muscle cells (= center of 5a). **C:** Reduced terms of gene ontology analysis of genes listed in Fig. 5b. **D:** List of genes from intersection of “cluster genes” for each differentiation (clusters 1 and 4 from Fig. 2j and Fig. 3ji and clusters 1 and 2 from Fig. 4j) with risk genes for non-syndromic orofacial clefting. **E:** List of genes from intersection of “cluster genes” for each differentiation (clusters 1 and 4 from Fig. 2j and Fig. 3j and clusters 1 and 2 from Fig. 4j) with orofacial cleft risk genes. **F:** Heatmap of hypoxia-attenuated OFC risk genes (= all genes from Fig. 5d,e). Scaled normalized counts for normoxic (21% O_2_) and hypoxic conditions (0.5% O_2_) for proliferating O9-1 cells (Pro) and all differentiations (CC = chondrocytes, OB = osteoblasts, SMC = smooth muscle cells) are shown. **G:** Gene ontology analysis of hypoxia-attenuated OFC risk genes (= all genes from Fig. 5d,e). **H:** STRING analysis of hypoxia-attenuated OFC risk genes (= all genes from Fig. 5d,e).

### Several OFC risk genes are poorly induced during hypoxic differentiation processes

For an evaluation of the clinical relevance of our findings, we compared the hypoxia-attenuated genes with known risk genes for OFC. First, we compared hypoxia-attenuated genes of chondrocyte, osteoblast and smooth muscle differentiation with risk genes for nsOFC (Fig. 5D) (Dixon, M. J. et al., 2011; Hoebel et al., 2017; Ishorst et al., 2022; Ruff et al., 2022; Thieme et al., 2021). We did not detect any nsOFC risk genes whose induction was compromised during hypoxic chondrocyte differentiation, but we found four corresponding to osteoblast differentiation and seven for smooth muscle differentiation. Interestingly, three of those nsOFC risk genes were shared between osteoblast and smooth muscle differentiation: Fibroblast growth factor receptor 2 (*Fgfr2*), Glutathione S-transferase theta-1 (*Gstt1*) and Thromboxane A2 receptor (*Tbxa2r*).

We extended our analyses to other OFC risk genes (i.e. syndromic OFC risk genes and genes validated in mutant mice) listed in the CleftGeneDB (https://bioinfo.uth.edu/CleftGeneDB/), in gene sets related to OFC (https://www.gsea-msigdb.org/) and in literature (Funato et al., 2015). We identified several known risk genes whose differentiation-dependent induction is compromised in hypoxia (Fig. 5E). These were 16 genes in chondrocytes, 15 in osteoblasts and 20 in smooth muscle cells. Intriguingly, we found three genes with impaired induction in all differentiation paradigms, namely the actin gene *Actg2* (Actin gamma-2, smooth muscle), the cell surface receptor *Boc* (Brother of Cdo) and the cysteine dioxygenase *Cdo1*. These three common genes again pointed to interference of hypoxia with cytoskeletal organization and amino acid metabolism during differentiation of CNCC.

In order to gain an overview of the expression of the identified OFC risk genes in the three differentiation processes, we performed clustering with hclust in R (Fig. 5F). Interestingly, many genes were not only induced in one differentiation paradigm, but also in others, even though to a lower extent. For example, *Fgfr2* is a hypoxia-attenuated gene during osteoblast differentiation, but also weakly induced in chondrocyte and smooth muscle differentiation. Furthermore, *Col11a1* and *Col11a2* were expectedly induced during chondrocyte differentiation and were hypoxia-attenuated, but also induced during the other differentiation paradigms. Notably, some of the smooth muscle-attenuated genes such as *Maf* and *Nbl1* were even stronger induced in chondrocyte or osteoblast differentiation and were not attenuated in hypoxia during these differentiation processes. We conclude that our identified hypoxia-attenuated genes often are not specifically induced in one differentiation process. Rather, they seem to be involved in other differentiation processes as well. Moreover, hypoxia-attenuated induction is often restricted to a specific differentiation process rather than being compromised by hypoxia in general.

We performed a GO term analysis of the OFC risk genes identified in our dataset and found terms related to our three differentiation paradigms (Fig. 5G): (1) chondrocytes (cartilage development, extracellular matrix organization), (2) osteoblasts (skeletal system development, ossification) and (3) smooth muscle cells (muscle organ development, ventricular cardiac muscle tissue morphogenesis, cell migration involved in heart development). Moreover, a large number of terms (eight out of thirty) was related to general development: animal organ development, multicellular organismal process, limb morphogenesis, regulation of cell differentiation, regulation of cell communication, embryonic pattern specification, developmental growth and cell fate commitment. Intriguingly, we detected two terms directly related to orofacial development, namely roof of mouth development and craniofacial suture morphogenesis. We therefore conclude that our approach yielded genes that are important during differentiation of neural crest-derived orofacial structures and whose expression is drastically reduced under hypoxic conditions.

To further disclose correlations between the hypoxia-attenuated genes, we performed a STRING analysis (https://string-db.org/, Version: 12.0) (Szklarczyk et al., 2023) of the hypoxia-attenuated OFC risk genes (Fig. 5H). Expectedly, the different collagen genes *Col2a1, Col9a1, Col9a2, Col11a1* and *Col11a2* clustered together, pointing to the impaired differentiation of CNCC to chondrocytes under hypoxic conditions. *Col2a1*, which was hypoxia-attenuated during chondrocyte and osteoblast differentiation, was related to upstream signaling molecules Bmp4 and Fgf18, consistent with the regulatory roles of Fgf and Bmp signaling in osteoblast and chondrocyte differentiation. The connected cluster included Dlx5, a transcription factor with a prominent role in osteoblast and chondrocyte differentiation that was also hypoxia-attenuated during osteoblast and chondrocyte differentiation. In general, half of the 34 identified hypoxia-attenuated OFC risk genes were part of this cluster, pointing to common mechanisms of differentiation that are compromised during hypoxia.

## DISCUSSION

Epidemiologic knowledge about OFC as the most frequent congenital malformation of orofacial structures is quite well established. Non-syndromic OFC are caused by an interplay of environmental factors and genetic risk factors (Dixon, Michael J. et al., 2011; Mossey et al., 2009). The genetic risk factors are being resolved by human geneticist studies (Ishorst et al., 2022; Leslie and Marazita, 2013; Ludwig et al., 2012). With a few exceptions, however, molecular explanations remain missing why a certain single nucleotide polymorphism might lead to orofacial clefting (Mangold et al., 2016; Rahimov et al., 2008). Also for environmental factors, in-depth knowledge about the molecular mechanisms that lead to OFC is scarce. This includes hypoxia during pregnancy. Experimental evidence for this relationship was already obtained in 1981 in animal studies (Millicovsky and Johnston, 1981), and clinical trials showed nearly 25 years ago an association of OFC with pregnancy in high altitudes (Castilla et al., 1999). Despite this long time, knowledge about the molecular mechanism behind hypoxia-induced OFC formation is still lacking. We here show for the first time that hypoxia severely affects the differentiation potential of CNCC to all three tested derivatives relevant for orofacial development, i.e. chondrocytes, osteoblasts and smooth muscle cells. In contrast, proliferation rates were affected only very mildly, if at all. Proliferating CNCC nevertheless sensed the drop in oxygen levels, as shown by qRT-PCR of typical hypoxia marker genes and as indicated by typical, hypoxia-related GO terms of induced genes identified by bulk RNA-seq. Of note, the central transcriptional regulator mediating transcriptional induction due to low oxygen levels, hypoxia-inducible factor-1alpha (Hif1a), is expressed in neural crest cells and is important for correct migration of neural crest cells to the heart region (Compernolle et al., 2003; Espina et al., 2018). We cannot exclude that in the in vivo situation proliferation of CNCC is also affected by low oxygen levels because nutrients might not be available in such excess as they are in cell culture. Nevertheless, Hif1a seems to act in an oxygen-independent manner on migrating neural crest cells (Compernolle et al., 2003; Espina et al., 2018). In fact, mild hypoxic conditions are beneficial for correct delamination of emigrating neural crest cells (Scully et al., 2016).

The effect of hypoxia on the differentiation potential of CNCC was dramatic: the differentiation-dependent induction of typical marker genes such as *Col2a1, Col11a2* (chondrocytes), *Msx1, Runx2* (osteoblasts) and *Acta2, Tagln* (smooth muscle cells) was not detectable or lowered dramatically to less than 20% of normal values. One might assume that differentiation of CNCC totally fails under hypoxic conditions. However, this is not the case, as GO terms of genes induced under hypoxic conditions were still related to the specific differentiation schemes. Moreover, although the majority of the 200 most upregulated genes in each differentiation paradigm showed drastically compromised induction, other genes were still robustly induced under hypoxic conditions. Thus, specific mechanisms seem to exist that allow induction of certain genes even under hypoxic conditions while induction of others is impaired. The key role of oxygen supply for correct differentiation of neural crest cells is underlined by the finding that hypoxia evokes dedifferentiation processes in neural crest-derived neuroblastoma cells in vitro and in vivo (Jögi et al., 2002).

A central cause for the reduced differentiation potential might be a compromised organization of the cytoskeleton and altered cell adhesion on the one hand, and disturbances of amino acid metabolism on the other. GO terms related to these processes were identified for genes whose induction was hampered in all three paradigms. Although the total number of identified genes with constrained induction in all differentiation processes is low, amino acid metabolism or cytoskeletal organization and cell adhesion may be worthwhile to investigate as targets for medication. Cytoskeletal organization and ECM-cell interaction are important for proper function of chondrocytes, osteoblasts and smooth muscle cells (Gao et al., 2014; Gould et al., 2021; Sweeney and Hammers, 2018). Furthermore, amino acid metabolism is also affected in CNCC in which subunits of the Ep400/Tip60 complex are deleted (Gehlen-Breitbach et al., 2023), pointing to the relevance of this metabolic pathway for craniofacial development. Mutations of Ep400/Tip60 complex subunits give rise to orofacial malformations in homozygous mouse mutants and to OFC in heterozygous mouse mutants and in patients (Gehlen-Breitbach et al., 2023; Humbert et al., 2020).

The relevance of our findings is highlighted by the fact that we detected several known risk genes for OFC among our genes with impaired hypoxic induction. Among them were such well-known candidate genes as *Fgfr2* (fibroblast growth factor receptor 2) (Slaney et al., 1996), *Bmp4* (bone morphogenetic protein 4) (Suzuki et al., 2009) and collagen genes (Brown et al., 1992; Melkoniemi et al., 2000). Nearly all of these 34 OFC risk genes were induced in each of our differentiation regimes. Nevertheless, induction rates varied between the different paradigms. This was in accord with the known roles of the genes in biological processes. Interestingly, not every induction of every gene was hampered by hypoxia. Rather, sometimes only a specific differentiation paradigm was affected by reduced induction. This points to expected differential mechanisms of gene induction for each differentiation process, but also to differential influences of hypoxia on these mechanisms on specific genes. GO terms of the 34 OFC risk genes with hypoxia-attenuated induction were related to the three differentiation regimes, to orofacial development and to general developmental processes, underlining the validity of our findings. STRING analysis identified interactions in a cluster for half of these genes, pointing to interconnected functions of the gene products.

Three OFC risk genes showed hypoxia-attenuated induction in each differentiation process, possibly pointing to their central roles. The fact that smooth muscle actin γ-2 (*Actg2*) is among them again points to the importance of cytoskeletal organization in this context. Despite its name, *Actg2* is not specifically expressed in smooth muscle cells. It is also expressed in osteoblasts and highly induced in early phases of osteoblast differentiation (Sebastian, 2016). Intriguingly, chondrocyte expression of *Actg2* is five times higher in neural crest-derived cells than in other chondrocytes (Hellingman et al., 2011). The second gene is the hedgehog co-receptor brother of CDO (*Boc*) (Tenzen et al., 2006). In addition to its long-known role in myoblast differentiation its function in neural crest cell migration was detected recently (Lencer et al., 2023). Finally, the cysteine dioxygenase *Cdo1* was among the identified commonly constrained genes. *Cdo1* is induced during osteoblastic differentiation of bone marrow stromal cells and serves as a negative regulator of osteogenic differentiation (Zhao et al., 2016). Moreover, it is expressed in chondrocytes and vascular smooth muscle cells (Forgan et al., 2017; Sun et al., 2024). The presumptive importance of amino acid metabolism for CNCC differentiation in the context of hypoxia is pointed out by the role of Cdo1 as a key enzyme in cysteine catabolism (Chen et al., 2023).

Of course, the drastically low oxygen levels we have chosen as an experimental paradigm (0.5%) do not equal the in vivo situation. Nevertheless, atmospheric oxygen levels drop to below 14% (equaling 2/3 of normal atmospheric oxygen) at altitudes above 3,300 m, which is the altitude where La Paz, Bolivia, is located. Atmospheric oxygen levels influence arterial oxygen saturation (SpO_2_) and mean SpO_2_ of healthy newborns drops to near 90% at this altitude (Li et al., 2023; Subhi et al., 2009). Naturally, the dramatic effects we observed in our cell culture model at 0.5% O_2_ would in vivo be detrimental for craniofacial development. The seemingly mild decrease to SpO_2_ levels of 90% may imply that this drop would not impact CNCC differentiation. It should be noted, however, that 90% SpO_2_ is the standard 2.5^th^ percentile that is considered as the cutoff for identifying hypoxemia and indicates the need for oxygen therapy (Li et al., 2023; Subhi et al., 2009). It is therefore highly conceivable that the effects detected by our experiments will translate into the in vivo situation.

## MATERIALS AND METHODS

### Cell culture

The murine cranial neural crest cell line O9-1 was purchased from Sigma-Aldrich and cultured essentially as described (Gehlen-Breitbach et al., 2023; Ishii et al., 2012). Briefly, cells were seeded on Matrigel-coated cell culture plates. Proliferation medium was STO- conditioned DMEM (Gibco) containing 15% fetal calf serum (FCS), 1% Penicillin/Streptomycin, 0.1 mM non-essential amino acids and 55 µM beta-mercaptoethanol to which 25 ng/ml bFGF and 5 ng/ml LIF had been added. Medium was changed every second day, passaging was performed with Accumax solution (Sigma-Aldrich) at a confluence of up to 75% with a reseeding density of 1:40. For normal atmospheric oxygen (21% O_2_) cells were kept in a CO_2_ incubator (Eppendorf). For hypoxic conditions (0.5% O_2_) cells were cultured in a hypoxic chamber (Whitley H35 HEPA) for at least 24 h.

When O9-1 cells had reached nearly 100% confluence, proliferation medium was changed to differentiation medium. Osteoblast differentiation medium was alpha-MEM (Gibco) containing 10% FCS, 1% Penicillin/Streptomycin, 10 mM beta-glycerophosphate, 0.1 µM dexamethasone and 100 ng/ml BMP-2. Cells were differentiated in this medium for 10 days, medium was changed every second day. For chondrogenic differentiation, cells were first cultured in osteoblast differentiation medium for three days, followed by micro mass culture on matrigel-coated plastic ware in chondrogenic differentiation medium for seven days. This medium consisted of alpha-MEM with 5% FCS, 1% Penicillin/Streptomycin, 1x Insulin-Transferrin-Selenium (Gibco), 0.1 µM dexamethasone, 10 ng/ml TGF-beta3 and 50 ng/ml BMP-2. Medium was changed every second day. Smooth muscle differentiation medium consisted of DMEM (Gibco) with 10% FCS and 1% Penicillin/Streptomycin. Smooth muscle differentiation was performed for three days, medium was changed after two days. For differentiations in hypoxic conditions, cells were proliferated at atmospheric oxygen levels. Differentiation was started in the hypoxic chamber with media preconditioned to 0.5% O_2_. Medium changes were also performed with medium that had been preconditioned to 0.5% O_2_.

### RNA extraction and qRT-PCR

RNA was extracted with the RNeasy plus kit (Qiagen) according to manufacturer’s instructions. After reverse transcription with SCRIPT cDNA synthesis kit (Jena bioscience), quantitative reverse transcriptase PCR (qRT-PCR) was performed on a Roche LightCycler 96 with the primer pairs listed in supplementary table S2. Quantification was performed by the ΔΔ-Ct method with Ct values normalized to the housekeeping gene *Rplp0*.

### RNA-seq and related analyses

RNAs of four biologically independent O9-1 samples which had been cultivated in proliferating as well as differentiating (chondrocyte, osteoblast, smooth muscle cell) conditions at 21% O_2_ as well as three respective biologically independent samples at 0.5% O_2_ were used for library generation (400ng per sample, Novogene NGS Stranded RNA Library Prep Set (PT044)), poly-A enrichment and RNA sequencing on the Illumina Sequencing PE150 platform with a read length of 2 × 150 bp and an output of ∼60 M reads per sample (Novogene, Cambridge). The generated sequence data were mapped onto the *Mus musculus* reference genome mm10 using hisat2 (version 2.0.5), assembled via Stringtie (version 1.3.3b) and quantified using featureCounts (version 1.5.0-p3) by the sequencing company. Bioinformatic analysis was performed using R version 4.3.0. The Bioconductor package DESeq2 (version 1.42.1) (Love et al., 2014) was applied for differential expression analysis with default parameters. The requirements for deregulation were an absolute value of the logarithmic fold change of at least 1 and a group counts’ mean larger than 100. The R packages clusterProfiler (version 4.10.1) (Wu et al., 2021), topGO (version 2.54.0) (Adrian Alexa, d 2023), GeneTonic (version 2.6.0) (Marini et al., 2021) and rrvgo (version 1.14.2) (Sayols, 2023) were used for gene ontology analysis, reduction of terms and visualization. The package STRINGdb (version 2.14.3) (Szklarczyk et al., 2023) was used for STRING analysis. The function hclust was used for hierarchical clustering of genes in heatmaps with default parameters.

### Cell counting and life/dead staining

For comparison of growth rates, O9-1 cells were re-seeded with progeny kept at 21% O_2_ or at 0.5% O_2_ in proliferation medium. After 24 h, 48 h and 72 h phase contrast photographs were taken on a Zeiss AxioVert A1 microscope. Cell numbers were counted with the Adobe Photoshop counting tool on at least three photographs from three independent experiments. Live/dead staining was performed with a Live/Dead Staining Kit (PromoCell) according to manufacturer’s instructions. Fluorescence photographs were taken on a Zeiss AxioVert A1 microscope. Quantification was performed with the Adobe Photoshop counting tool on at least three fluorescence photographs from three independent experiments.

## DATA AVAILABILITY

All data are provided in the paper and its supplement or have been submitted to GEO under accession number GSE273808.

## ACKNOWLEDGEMENTS

TS was financed by grants from the Interdisciplinary Center for Clinical Research (IZKF) at the University Hospital of the University of Erlangen-Nuremberg to LG and MiW (grant number E28) and to MaW (grant number P140). Furthermore, this project received funding by the Deutsche Gesellschaft für Kieferorthopädie to MaW (grant title “Identifizierung von molekularen Ursachen hypoxiebedingter orofazialer Spalten”).

## AUTHOR CONTRIBUTIONS

MaW, TS and GR conceived the study and designed experiments. GR and TS performed experiments. TS and MaW performed statistical analysis and interpreted the experiments with the help of AK. LG funded the project as well. LG and MiW helped supervise the project. MaW, TS and AK wrote the manuscript with the help of LG and MiW.

## SUPPLEMENTARY FILES

**Table S1:** Lists of genes whose expression is constrained during differentiation of O9-1 cells to chondrocytes (CC), osteoblasts (OB) or smooth muscle cells (SMC)

**Table S2:** Sequences of PCR primers used in this study

