## Supplementary material for "Hypoxia impedes differentiation of cranial neural crest cells into derivatives relevant for craniofacial development": Supplemenal Table 1

| CC | OB | SMC |
|---|---|---|
| 4833422C13Rik | 1500015O10Rik | 4930474H06Rik |
| Actg2 | Abat | Abca9 |
| Adamts16 | Abca9 | Ackr4 |
| Adamtsl1 | Abi3bp | Acot11 |
| Adamtsl2 | Ackr4 | Acta1 |
| Agt | Actg2 | Acta2 |
| Alpl | Adamtsl1 | Actg2 |
| Alx4 | Adcy5 | Actn3 |
| Arhgef6 | Agt | Adamtsl1 |
| Aspn | Aldh1a1 | Adh1 |
| Bmp8a | Aldh1a7 | Angpt1 |
| Boc | Aoc3 | Anpep |
| C130050O18Rik | Apod | Aoc3 |
| C1qtnf3 | Arhgef6 | Apod |
| Cacna1g | Art3 | Arhgap28 |
| Cacna1h | Aspn | Aspn |
| Capn6 | Bmf | Atp9a |
| Cdh6 | Boc | Bdh2 |
| Cdo1 | C1s1 | Bmf |
| Cfh | Cacna1g | Bmp3 |
| Chrdl1 | Cacna1h | Bmp4 |
| Cnr1 | Cacnb4 | Boc |
| Col10a1 | Capn6 | C1ra |
| Col11a2 | Ccdc3 | C1s1 |
| Col27a1 | Cdh18 | Cacna1g |
| Col2a1 | Cdo1 | Cacnb4 |
| Col9a1 | Col11a1 | Calml4 |
| Col9a2 | Col12a1 | Capn6 |
| Colgalt2 | Col23a1 | Casq2 |
| Comp | Col2a1 | Ccdc3 |
| Cpm | Col8a2 | Cdh11 |
| Cybrd1 | Col9a1 | Cdo1 |
| Dlx5 | Col9a2 | Cfh |
| Dpep1 | Cpm | Chn2 |
| Ednra | Dlx5 | Chrdl1 |
| Eln | Dpep1 | Chst2 |
| Enpp2 | Dpyd | Cib2 |
| Erg | Ecm2 | Cldn19 |
| F13a1 | Egfl6 | Clec11a |
| Fam180a | Ehd3 | Clu |
| Fam189a2 | Eln | Cnn1 |
| Fam20a | Erg | Col10a1 |
| Fgfr3 | Fam107a | Col11a1 |
| Fibin | Fam180a | Col12a1 |
| Fmo1 | Fbln7 | Col23a1 |
| Frzb | Fbxo2 | Col27a1 |
| G0s2 | Fgf18 | Col8a2 |
| Gas6 | Fgfr2 | Cp |
| Gldn | Fgfr3 | Cped1 |

| | | |
|---|---|---|
| Gm26778 | Fhl5 | Cpxm1 |
| Gm27483 | Fibin | Ctsf |
| Gm27786 | Fmo1 | Dact1 |
| Gpbar1 | Fmo2 | Dcn |
| Gprc5c | Frzb | Dusp27 |
| Gprin3 | Fxyd1 | E030013I19Rik |
| Gpx3 | Galnt9 | Ecm2 |
| H19 | Gas6 | Ednra |
| Hapln1 | Gdpd2 | Eepd1 |
| Hey1 | Gm15655 | Egfl6 |
| Hif3a | Gm26776 | Enpp5 |
| Hp | Gm26778 | Epha3 |
| Hpgd | Gm42778 | Erg |
| Ibsp | Gpr157 | Fabp3 |
| Igfbp5 | Gpr88 | Fam180a |
| Islr | Gstt1 | Fat4 |
| Kcnh1 | Hapln1 | Fbln7 |
| Kcnj15 | Hif3a | Fbxl7 |
| Kctd12 | Hmgcs2 | Fgfr2 |
| Kif26b | Ifitm10 | Fgl2 |
| Klhl33 | Igdcc4 | Flrt1 |
| Lama4 | Igf1 | Fmo1 |
| Lbp | Islr | Fmod |
| Limch1 | Itga1 | Frzb |
| Lrp4 | Kcnmb4 | Fxyd1 |
| Lum | Kctd12 | Gdpd2 |
| Map2 | Kif26b | Gm15655 |
| Map7d2 | Lama2 | Gm27483 |
| Matn4 | Lama4 | Gm27786 |
| Mdfi | Limch1 | Gper1 |
| Mdga1 | Lmod1 | Gpx3 |
| Megf10 | Lrrc75b | Gstt1 |
| Mepe | Lum | H19 |
| Mical1 | Mchr1 | Hmcn1 |
| Moxd1 | Megf6 | Hr |
| Nrep | Mest | Igdcc4 |
| Ostn | Mme | Igf1 |
| Panx3 | Mmp15 | Igsf10 |
| Papss2 | Mrvi1 | Il17rd |
| Parm1 | Myl9 | Islr |
| Pdgfrl | Myom1 | Itga1 |
| Ptgis | Ndst4 | Itga11 |
| Pth1r | Negr1 | Itga8 |
| Ptpru | Nkd1 | Kctd12 |
| Reln | Nol4l | Kif26b |
| Robo2 | Ogn | Lama4 |
| Rtl3 | Otulinl | Ldb2 |
| Rxfp3 | Pamr1 | Lgr6 |
| Scin | Pappa | Limch1 |
| Scube1 | Pcp4l1 | Lims2 |

| | | |
|---|---|---|
| Serpina3m | Pdgfrl | Lmod1 |
| Shank1 | Pgm5 | Lpl |
| Slc2a10 | Plxdc2 | Lrp4 |
| Slc43a2 | Plxnc1 | Lrrc17 |
| Slc7a2 | Postn | Lrrc75b |
| Sparcl1 | Prlr | Maf |
| Srpx | Prss12 | March3 |
| Tll1 | Ptgis | Megf6 |
| Tmem154 | Pth1r | Mest |
| Tmem229b | Ramp1 | Mfap2 |
| Tmie | Rassf4 | Mfap4 |
| Tmtc1 | Reps2 | Mgll |
| Tspan18 | Robo2 | Mical1 |
| Vit | Rtl3 | Mme |
| Wif1 | Rtl8c | Mn1 |
| Wscd2 | Rtn4rl1 | Mrvi1 |
| | Scara5 | Myl9 |
| | Selenop | Mylk |
| | Sema3g | Myom1 |
| | Shank1 | Nbl1 |
| | Slc43a2 | Nid2 |
| | Slc7a2 | Nkd1 |
| | Smoc2 | Nrep |
| | Sncaip | Ogn |
| | Stk32b | Olfml2a |
| | Sult1a1 | Otulinl |
| | Susd5 | Pappa2 |
| | Tbxa2r | Pdgfrl |
| | Tmtc1 | Pdk2 |
| | Tnn | Phactr1 |
| | Tox | Plxdc2 |
| | Tril | Plxnc1 |
| | Tspan18 | Pnmal2 |
| | Wdr72 | Podn |
| | Wscd2 | Postn |
| | Zfp185 | Prelp |
| | | Prr5l |
| | | Prss12 |
| | | Prss35 |
| | | Ptgis |
| | | Rab3il1 |
| | | Ramp1 |
| | | Rasl11b |
| | | Reps2 |
| | | Rgs4 |
| | | Rnf144a |
| | | Robo2 |
| | | Rspo1 |
| | | Rtn4rl1 |
| | | Selenop |

Sema3g
Sema6c
Serpine2
Sfrp1
Sgk3
Shank1
Slc24a3
Slc40a1
Slc7a2
Smarca1
Smoc2
Snap91
Sntb1
Socs2
Sorcs1
Sparcl1
Spats2l
Srpx
Svep1
Tagln
Tbxa2r
Tcn2
Tent5a
Tll1
Tmem140
Tmtc1
Tnn
Tox
Wisp2
Ypel2
