## Supplementary material for "Hypoxia impedes differentiation of cranial neural crest cells into derivatives relevant for craniofacial development": Supplemenal Table 2

| target gene | 5' primer | 3' primer |
|---|---|---|
| Acta2 | CACCATGTACCCAGGCATTG | CTGGAAGGTAGACAGCGAAG |
| Col11a2 | CTTCCGGGTGTTCTGCAAC | CCAGAGCAAGGGTAGGAGAC |
| Col2a1 | AAGAACAGCATCGCCTACCT | GGAGGTCTTCTGTGATCGGTR |
| Msx1 | CCAGAAGATGCTCTGGTGAAG | TTGGTCTTGTGCTTGCGTAG |
| Rplp0 | GACTGAGTACACCTTCCCACT | AGGCTGACTTGGTTGCTTTG |
| Runx2 | CCTGAACTCTGCACCAAGTC | GAGGTGGCAGTGTCATCATC |
| Tagln | CAACAAGGGTCCATCCTACGG | ATCTGGGCGGCCTACATCA |
